# Landscape and functional repertoires of long noncoding RNAs in the pan-cancer tumor microenvironment using single-nucleus total RNA sequencing

**DOI:** 10.1101/2023.12.03.569806

**Authors:** Tongqiang Fan, Shengyu Ni, Haide Chen, Ziye Xu, Longjiang Fan, Yongcheng Wang

**Author notes:** Corresponding Authors: Yongcheng Wang; Longjiang Fan.

## Abstract

Intratumor heterogeneity (ITH) plays crucial roles in tumor progression. However, the atlas of long noncoding RNAs (lncRNAs) in the context of ITH across multiple cancer types remains largely unexplored. Here, we analyze over 800,000 cells from ten different cancer types generated from the random-primed single-nucleus total RNA sequencing and provide a systematic landscape of lncRNAs in tumor microenvironment (TME) and malignant programs. Our study employe a robust cell annotation pipeline called scAnnotation, which allows us to identify 39 distinct cell types within the pan-cancer TME. By applying stringent criteria, we identify thousands of reliable marker genes, including both mRNAs and lncRNAs. Next, we identify sets of cell type-specific lncRNA-mRNA pairs by our LncPairs algorithm. Moreover, we identify nine expression meta-programs (MPs) associated with diverse biological processes in malignant cells across multiple cancer types. MP-specific lncRNA-transcription factor (TF) regulatory networks are further constructed and key lncRNAs and regulons that exert control over MP-specific gene expression are identified. The comprehensive atlas of lncRNAs in the pan-cancer context, coupled with the bioinformatics tools tailored for the random-primed datasets, is expected to accelerate advancements in the field of lncRNA research at the single-cell resolution.

## Introduction

Intratumor heterogeneity (ITH) is a well-established phenomenon in cancer biology that contributes to tumor progression, metastasis, and resistance to therapy^1,2^. Single-cell/nucleus RNA sequencing (sc/snRNA-seq) technologies have revolutionized our understanding of ITH and reveal a remarkable degree of heterogeneity within individual tumors^3–7^. Previous studies in melanoma^3,4^, glioblastoma (GBM)^5^, head and neck squamous cell carcinoma (HNSCC)^6^, and pancreatic ductal adenocarcinoma (PDAC)^7^ have identified ITH expression programs, such as skin-pigmentation, epithelial-mesenchymal transition (EMT), and epithelial senescence, with important functional implications. While malignant cells exhibit high heterogeneity across different cancer types and tissues, the presence of similar ITH programs in multiple cancer types enables pan-cancer studies of malignant cells^8,9^. Recently, a comprehensive pan-cancer study systematically investigates dozens of ITH programs and defines them as meta-programs (MPs)^8^. These MPs provide a framework for understanding the common features and functional implications of ITH across multiple cancer types.

Long noncoding RNAs (lncRNAs) are a class of non-coding RNAs that are longer than 200 nucleotides and exhibit spatiotemporal and tissue-specific expression patterns, making them potential cell type marker genes. Recent advancements in high-throughput sequencing technologies, particularly single-cell sequencing, have greatly expanded our knowledge of lncRNA genes in various mammalian tissues and cells^10,11^. Cumulative evidences have revealed that lncRNAs exhibit significant cell heterogeneity and play critical roles in both immunity and tumor progression^12–15^. For instance, *H19*, *lnc13*, and *HOXA-AS2* have been found to be expressed in human and mouse immune cells, suggesting their role in regulating immune targets^12–14^. On the other hand, oncogenic and cancer-promoting lncRNAs, including *NEAT1*, *HERES*, *MALAT1*, and *HOTAIR*, have been reported to modulate tumorigenesis and cancer progression^15^. However, current lncRNA-associated single-cell research mainly rely on datasets generated from oligo-dT based single cell 3’ RNA sequencing technologies such as 10X Genomics Chromium platform and BD Rhapsody platform. These methods rely on the hybridization of barcoded oligo-dT primers to the poly(A) sequences of polyadenylated transcripts for RNA capture and complementary DNA (cDNA) synthesis, which suffer from low capture efficiency for lncRNAs^16^. Nevertheless, these limitations can be overcome using random-primed single-cell/nucleus total RNA sequencing technologies such as MATQ-seq, VASA-seq, snRandom-seq, and snHH-seq, which offer high sensitivity in detecting lncRNAs^17–20^.

In the field of cancer research, generalizing findings across different cancer types presents a significant challenge. As a result, the generated data sets exhibit inconsistent granularity and nomenclature, making cross-study comparisons and integrative analyses difficult. Moreover, in the realm of random-primed sequencing technology, there is a lack of systematic investigation into lncRNAs and corresponding analysis methods. While substantial progresses have been made in understanding protein-coding genes using these sequencing technologies, the same level of attentions and explorations have not been extended to lncRNAs. Consequently, our knowledge of the functional implications and regulatory mechanisms of lncRNAs in the context of random-primed sequencing remains limited. To address these challenges, it is imperative to establish standardized annotation systems that can facilitate data harmonization and enable more robust comparisons across studies. Additionally, concerted efforts are needed to advance the systematic investigation of lncRNAs within the framework of random-primed sequencing technologies.

In this study, we leverage an unprecedentedly extensive random-primed single-nucleus transcriptome dataset comprising over 800,000 cells derived from diverse tissues representing ten distinct cancer types. Through the application of the analysis pipeline and bioinformatics toolbox by this study, we meticulously construct an updated cell atlas of the pan-cancer microenvironment. Additionally, we identify and curate novel cell markers specifically tailored for the context of random-primed snRNA-seq technologies. A pivotal aspect of our investigation involves the comprehensive annotation of the tumor microenvironment (TME) lncRNA transcriptome. We specifically focus on unraveling the functional roles associated with the transcriptional heterogeneity of multiple cellular subpopulations within the TME. By shedding light on intricate transcriptional dynamics of the TME, our study aims to provide a foundational and invaluable resource for cell annotation in the domain of random-primed single-cell RNA sequencing technologies. Furthermore, our findings hold promise for guiding future explorations into the regulatory mechanisms of lncRNAs within the context of ITH.

## Result

### An updated cell atlas of pan-cancer microenvironment

In our previous studies^19–21^, we developed a random-primed single nucleus total RNA sequencing method for FFPE and frozen tissues, and applied it to profile 32 samples derived from nine distinct cancer types^20^ and 17 samples obtained from gliomas patients^21^, resulting in a combined dataset encompassing a total of 801,879 high-quality single-nucleus full-length total transcriptome data originating from 49 patients representing ten different cancer types. Leveraging this extended dataset, we construct an updated and refined transcriptional atlas of pan-cancer microenvironment (Fig. 1, Supplementary Fig. 1).

**Fig. 1.**
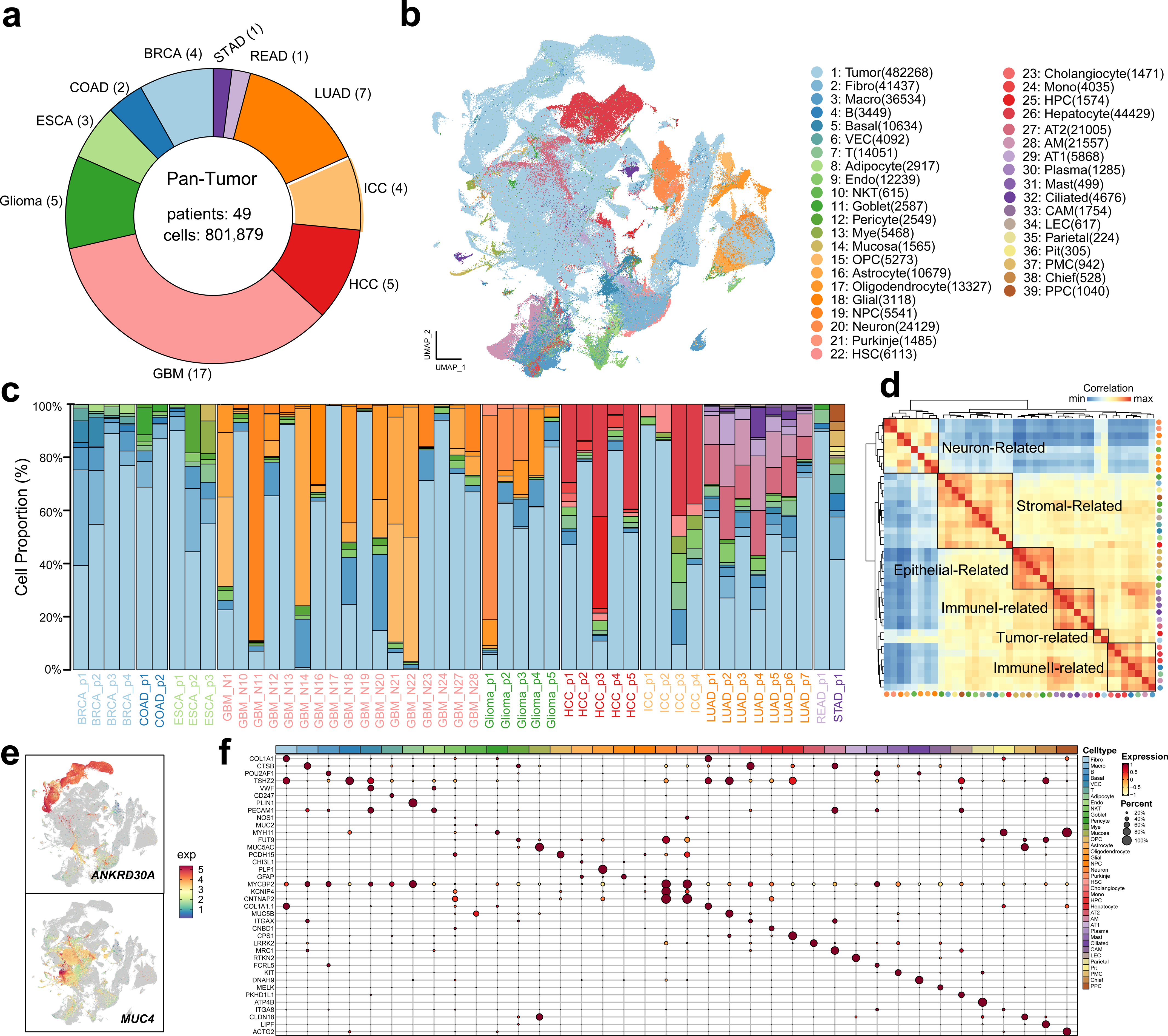
An updated tumor microenvironment using random-primed snRNA sequencing data. **a** Circle plot shows the samples used in this study. **b** UMAP clustering of 801,879 single cells from pan-cancer samples grouped by cell types. **c** Bar plot displaying the distribution of cells in each cluster based on the patient of origin. **d** Unsupervised hierarchical clustering heatmap shows transcriptome similarity of the cell clusters. Top 1000 variable features in each cluster were used for calculating the similarities. These clusters can be classified into six major clusters, neuron-related, stromal-related, epithelial-related, immune 1-related, tumor-related and immune 2-related. **e** UMAP plots showing the expression of two selected markers (*ANKRD20A* and *MUC4*) in clusters (i.e., cancer types). The rest were shown in Supplementary Fig. 1f. **f** Dot plot showing top one markers in each cluster.

In order to address the need for harmonizing annotations and identifying commonalities across different cancer types, we put forth a hypothesis that the integration of cells from diverse cancer types into a unified reference dataset could offer valuable insights. To achieve this objective, we employe a novel pipeline, termed as scAnnotation, which follows a robust framework encompassing comprehensive integration analysis of cell type annotation. This pipeline combines the strengths of both dataset-based and marker-based algorithms (See Methods, Supplementary Fig. 1d). By leveraging the scAnnotation pipeline, we successfully identify and characterize common cellular phenotypes across multiple studies and technologies. This approach facilitates the merging of disparate datasets, enabling joint analyses and enhancing our ability to discern shared features and patterns within the integrated dataset. Through this innovative methodology, we are able to effectively harmonize annotations and establish a reliable foundation for further investigations into the cellular landscape of various cancer types.

In this study, we analyze the 801,879 single-nucleus transcriptomes and visualize them in a two-dimensional space using the uniform manifest approximation and projection (UMAP) algorithm. We then apply scAnnotation to annotate the resulting clusters (Fig. 1b). Finally, we manually curate cell types, utilizing reference datasets and markers, and identify 39 major cell types, such as tumor cells, fibroblasts, macrophages, B cells, T cells, etc. Most of the cell types have a median of 4,035 cells per type, with the largest cell type being tumor cells (482,268 cells) and the smallest being parietal cells (224 cells). However, additional heterogeneity is observed, leading us to adopt an iterative strategy. We perform Louvain clustering on each major cell type, and observe that highly specific known marker genes are consistent with our annotation of the major cell types (Fig. 1, Supplementary Fig. 1f, Supplementary Data 1). For instance, the three most abundant cell types in the pan-cancer cell map are cluster 1 (tumor cells), cluster 26 (hepatocytes), and cluster 2 (fibroblasts). These three cell types are characterized by high expression of *MUC4*, *CPS1*, and *COL1A1*, respectively. Moreover, we identify various cell states, such as goblet cells (2,587 cells), mucosa (1,565 cells), cholangiocyte (1,471 cells), AT1 cells (5,868 cells), AT2 cells (21,005 cells), and ciliated cells (4,676 cells) among the large number of epithelial cells (Fig. 1b). Our annotation approach is robust as all cell types are present in multiple cancer types and patients (Supplementary Fig. 1b-c). The tumor cell type accounts for more than 60% of the total cells, while hepatocyte and fibroblast cells constate 5.5% and 5.2% of the total cells, respectively. Smaller clusters are also readily annotated. For example, cluster 6 (VEC, vascular endothelial cell) expresses *VWF* specifically, whereas cluster 31 (mast cell) expresses *KIT* specifically. We observe that some markers are expressed in a substantial proportion of cells in many clusters but much more highly expressed in one cluster (for example, *MYCBP2* in NPC, neural progenitor cell). Annotation of the basal cell clusters is more challenging due to the smaller number of known markers (for example, *ATP4B* in parietal cell, *MYH11* in pericyte cell).

Based on the comprehensive annotation of the pan-cancer cell atlas described above, we observe significant intra– and inter-tumor heterogeneity in the cellular composition of the TME, as well as distinct expression signatures specific to individual patients (Fig. 1), which are consistent with previous studies^22–24^. We further characterize the broach cell type within the TME by analyzing covariance patterns (Fig. 1d). Specifically, by examining the mean expression and co-expression of highly variable genes, we identify prominent covariance effects that contributed to defining the TME clusters. To gain a deeper understanding of the relationships between these clusters, we conduct pairwise cluster similarity analyses. This allows us to classify 39 major clusters into six meta-clusters, namely neuron-related, stromal-related, epithelial-related, immune I-related and immune II-related clusters. Importantly, even after equalizing the mean gene expression across clusters, we observe substantial differences between these meta-clusters, underscoring their distinct molecular characteristics and functional roles within the TME.

Together, by leveraging the power of random-primed snRNA-seq technologies and an integrative pipeline of cell type annotation (scAnnotation), we have generated a more comprehensive pan-cancer cell atlas. This resource provides unprecedented insights into the cellular composition and heterogeneity of the tumor microenvironment across a wide range of cancer types. Importantly, our approach enables investigation of lncRNAs at single-nucleus resolution, which may shed new light on their functional roles in cancer biology. We believe that this atlas will serve as a valuable resource for future research efforts aimed at understanding the complex interplay between different cell types within the TME, as well as the molecular mechanisms underlying cancer progression and metastasis. Overall, our study represents a significant step forward in our understanding of the cellular and molecular landscape of cancer and provides a foundation for future investigation in this field.

### Identification of lncRNA markers and lncRNA-mRNA pairs

In order to gain a deeper understanding of the data generated through random-primed technologies, we devise a rigorous workflow aimed at identifying cell type markers, as outlined in the Methods section (Fig. 2a). Through this meticulous screening process, we successfully identify 2,083 confident markers, all of which are protein-coding genes, representing 30 distinct cell types. The number of markers per cell type varies, ranging from 14 to 244 (Fig. 2b, Supplementary Data 2). Notably, we find that a significant majority (90.7%) of these markers, could be validate and corroborated through public databases (Supplementary Fig. 1e). For instance, within cluster 2, which represents fibroblast cells, we observe the highest expression levels of *COL1A1*, *COL1A2*, *COL3A1*, and *COL6A3*. These genes are well-known markers associated with fibroblast cells according to previous literature^24^. However, it is worth mentioning that certain genes had not previously been characterized as marker for their respective cell types. For example, our pipeline detects the presence of *ADAM12*, *LAMA2*, *SLIT3* and *HPSE2* as markers of cluster 2. These genes have not previously been described as markers of fibroblast cells. The results demonstrate the power and potential of our pipeline in uncovering novel markers that may have been overlooked in previous studies, further enhancing our understanding of the cellular landscape and heterogeneity within fibroblast populations.

**Fig. 2.**
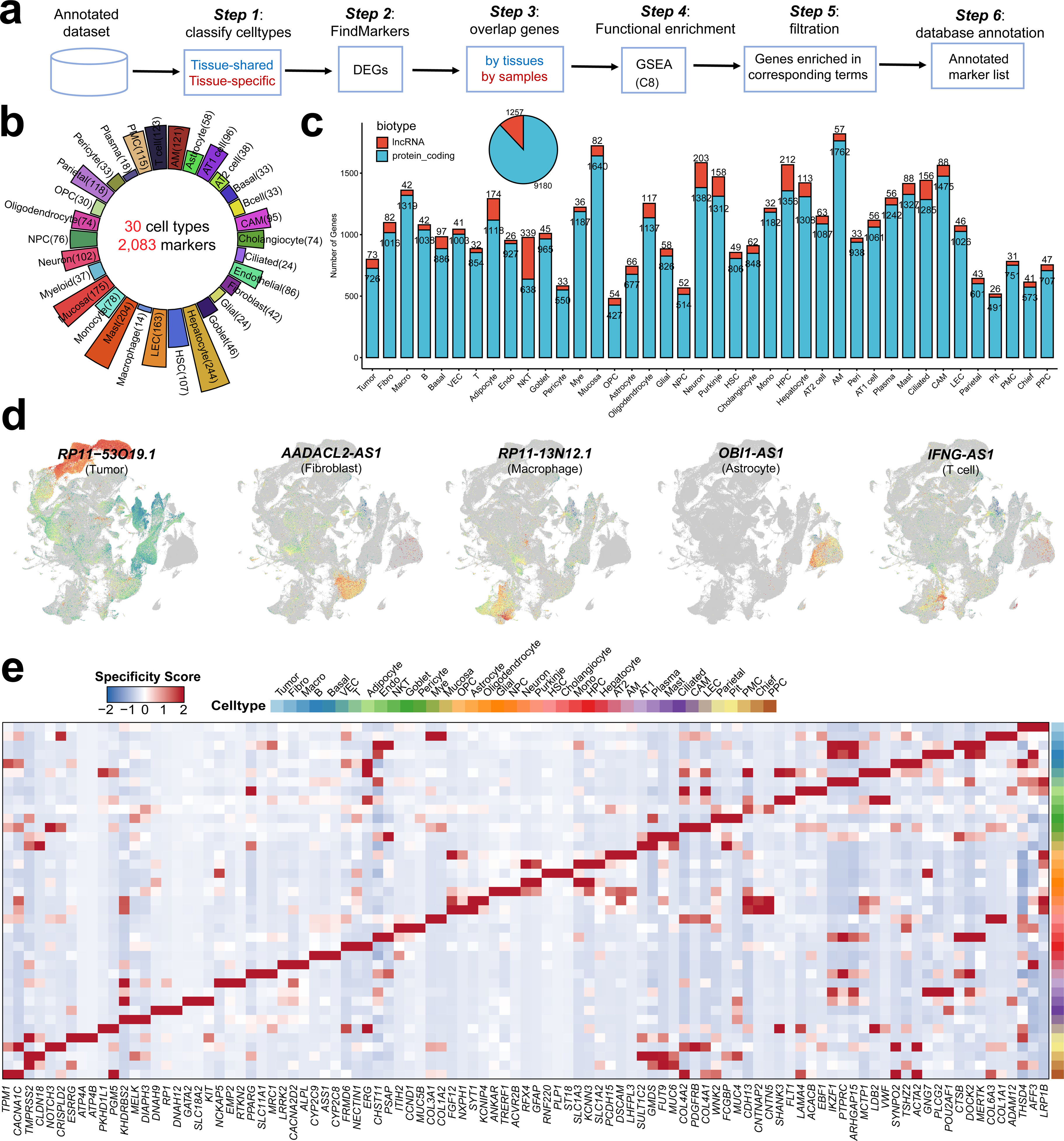
Identification of mRNA and lncRNA markers. **a** The pipeline to identify novel markers for random-primed snRNA-seq protocol. **b** Circled bar plot showing the number of markers of each cell type. **c** The number of mRNA and lncRNA markers in each cell type. Pie plot showing the total number of lncRNAs and mRNAs and stacked bar plot showing the number of markers in each cell type. **d** Expression profiles of selected lncRNA markers for each cell type. The rest of the markers were shown in Supplementary Fig. 2c. **e** Heatmaps of top three mRNA markers for each cell type based on their cell type specificity scores.

To gain insights into the role of lncRNAs in the pan-cancer TME, we conduct a comprehensive analysis to identify lncRNA markers associated with different cell types. Differential expression and correlation analyses are performed (See Methods). Across all major cell types, we observe a remarkable enrichment of cell-lineage specific expression for lncRNAs compared to protein coding genes. Specifically, out of the 11,129 lncRNA genes examined, 1,257 (11.3%) exhibits significant differential expression (adjusted *P* value < 0.01). On average, each cell type shows significant differential expression for 105 lncRNAs, with a range from 26 to 339 (Fig. 2c). To further investigate the potential interactions between lncRNAs and protein coding genes, we perform Pearson correlation analysis between the 1,257 differentially expressed lncRNAs and 2,083 confidential protein coding markers. Surprisingly, we identify 1,220 lncRNAs that exhibited significant correlations (correlation coefficient > 0.7, *P* value < 0.01) with the protein coding markers (Supplementary Fig. 2a). Notably, a substantial proportion of these correlated lncRNAs are found to be highly enriched in neuron-associated cell clusters, suggesting their potential involvement of neuron-related processes within the TME (Fig. 2c). The results warrant further investigation to elucidate the specific roles of these lncRNAs in the context of neuronal regulation and cancer progression.

In order to investigate the expression patterns of lncRNA markers within distinct cell types of the pan-cancer TME, we employe visualization methods utilizing UMAP embeddings (Fig. 2d, Supplementary Fig. 2c). Our analysis reveals highly specific expression patterns of corresponding lncRNAs within each cell cluster. For instance, we observe that *RP11-53O19.1*, a lncRNA previously reported to be overexpressed in breast cancer cells^25^, exhibits specific expression exclusively within tumor cells. This finding confirms and reinforces the association between *RP11-53O19.1* and breast cancer cells, further validating its potential as a distinctive marker for this cell type (Fig. 2d). Furthermore, our analysis identifies *OBI1-AS1* as a super-exclusive marker for astrocytes, corroborating previous findings reported by Mamivand et al^26^. The specific expression of *OBIA-AS1* within astrocytes provides further evidence of its roles as a highly specific marker for this particular cell type (Fig. 2d). Moreover, our investigation leads to the identification of *AADACL2-AS1* as a novel marker for fibroblasts and *RP11-13N12.1* as a marker for macrophages. These lncRNAs have not been previously recognized as markers for these respective cell types. This finding expands our knowledge of the molecular signatures associated with fibroblasts and macrophages, contributing to a more comprehensive understanding of their roles within the TME.

To identify cell type-specific lncRNAs and further elucidate the potential function of lncRNAs in pan-cancer TME, we conduct an analysis of co-expression profiles between mRNAs and lncRNAs (Fig. 2e, Supplementary Fig. 2b). The specificity scores exhibited by the identified lncRNAs are found to be comparable to those of mRNAs. Among the top 117 specific lncRNAs (the three most specific lncRNAs in each of the 39 clusters), the majority (111) are successfully matched to their annotations in Ensembl. This indicates the reliability and validity of our pipeline in accurately identifying known lncRNA markers. Notably, *TRHDE-AS1*, with significantly higher mean expression in adipocyte cells compared to other cell types, demonstrates the highest degree of cell type-specific enrichment among all genes analyzed (Supplementary Fig. 2b). This finding aligns with previous studies associating *TRHDE-AS1* with human adipocyte tissue development^27^. However, our analysis also reveals three lncRNAs, namely *LINC01060*, *RP11-231C18.3*, and *RP11-1017G21.4*, which exhibited significantly high specificity scores in basal cells, NKT (natural killer T) cells, and mucosal cells, respectively. These lncRNAs have not been previously reported to be correlated with their respective cell types. Their identification as potential novel markers underscores the capability of our marker identification pipeline to not only detect known markers but also uncover previously unrecognized associations between lncRNAs and specific cell types. These observations indicate the potential of our approach to aid in the discovery of novel markers, both in conjunction with mRNAs and lncRNAs, thereby expanding our understanding of the diverse cellular landscape and functional heterogeneity within the pan-cancer TME.

Previous investigations on lncRNA regulation have predominantly relied on bulk RNA sequencing data, which may overlook the intricate heterogeneity underlying lncRNA regulation at the individual cell level. To address this limitation and delve into the realm of cell-specific lncRNA-mRNA interactions, we develop a computational tool called LncPairs (Fig. 3a). By harnessing the capabilities of LncPairs, we successfully unveil a multitude of cell type and cancer type-specific lncRNA-mRNA pairs. Specifically, we identify 10,464 significant lncRNA-mRNA pairs (*P* value < 0.01) that exhibited cell type specificity, while an impressive 54,684 lncRNA-mRNA pairs demonstrate specificity within various cancer types (Fig. 3b-c). Notably, among the diverse cancer types analyzed, datasets from GBM exhibit the highest number of lncRNA-mRNAs pairs, warranting further exploration to unravel it unique molecular landscape. Within our comprehensive analysis, we discover numerous novel lncRNA-mRNA interactions that shed light on the intricate regulatory networks at play. For instance, our findings unveil a previously unknown interaction between the lncRNA *TRHDE-AS1* and the adipocyte-specific gene *LEP*, resulting in the formation of the cell-specific TRHDE-AS1-LEP pair. Furthermore, we observe an endothelial-enriched lncRNA (*LINC00607*)^28^, which interacts with three macrophage-specific genes (*CCL18*, *VSIG4* and *IL1A*)^29–31^ to form three macrophage-specific lncRNA-mRNA pairs: LINC00607-CCL18, LINC00607-VSIG4, and LINC00607-IL1A. These observations highlight the intriguing phenomenon whereby the ectopic expression of lncRNAs and mRNAs can form cell-specific lncRNA-mRNA pairs. Collectively, our findings provide compelling evidence that lncRNAs and their corresponding lncRNA-mRNA pairs play a crucial role in shaping and determining the cellular states. The identification and characterization of these interactions pave the way for a deeper understanding of the intricate mechanisms governing cell fate determination.

**Fig. 3.**
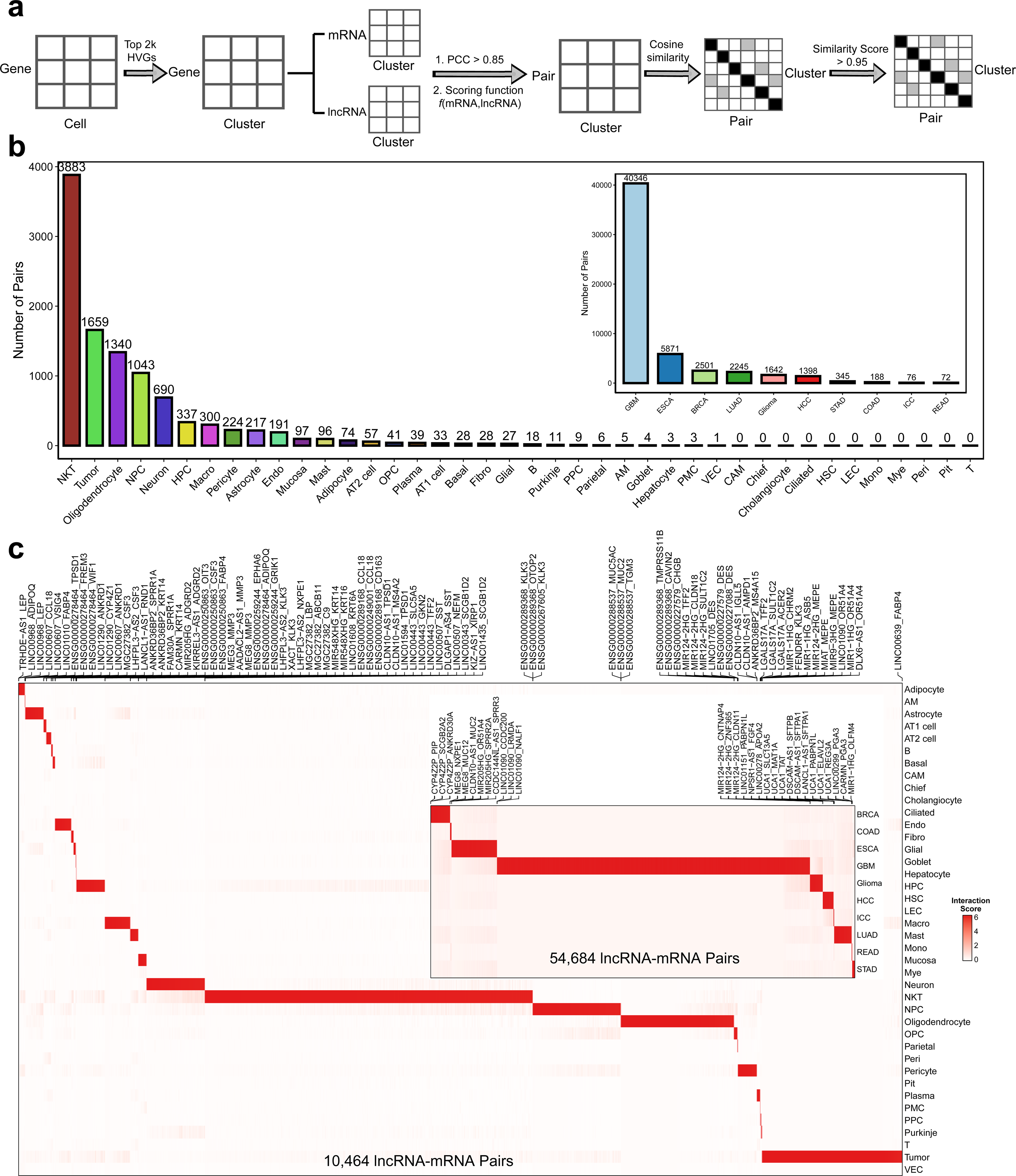
Identification of lncRNA-mRNA pairs at single-nucleus level. **a** The LncPairs algorithm for identification of cell/sample-specific lncRNA-mRNA pairs. **b** The number of pairs specific in each cell types (outer) and cancer types (inner). **c** Heatmap of the cell-specific lncRNA-mRNA pairs between cell types (outer) and cancer types (inner).

### LncRNA-associated malignant cell heterogeneity

To investigate the molecular underpinnings of malignant cells, we utilize the concept of MPs (Supplementary Data 4) and NMF to identify gene modules and assess the impact of lncRNAs on specific malignant cell states (See Methods). Firstly, we conduct a direct integration of datasets using our in-house pipeline and categorized malignant cell into distinct ITH programs (Supplementary Fig. 3a-b). Notably, following the filtration of programs associated with low quality or doublets, we curate a catalog comprising nine consensus modules, encompassing an impressive cell range from 1,476 to 185,874, which were then visually represented using UMAP (Fig. 4a). Remarkably, our analysis reveals that 77.8% (7 out of 9) of the identified ITH programs are recurrent patterns derived from multiple cancer types, underscoring the pervasive nature of these patterns across diverse malignancies. It is important to note that the primary expression data exhibited prominent batch effects, stemming from technical variations across different samples and tumor types, leading to the delineation of discrete clusters (Supplementary Fig. 3c). Further in-depth investigation uncovers that these distinct programs are enriched in specific organ systems, with a notable emphasis on the brain, while others extend across multiple organ systems and histologies (Fig. 4a).

**Fig. 4.**
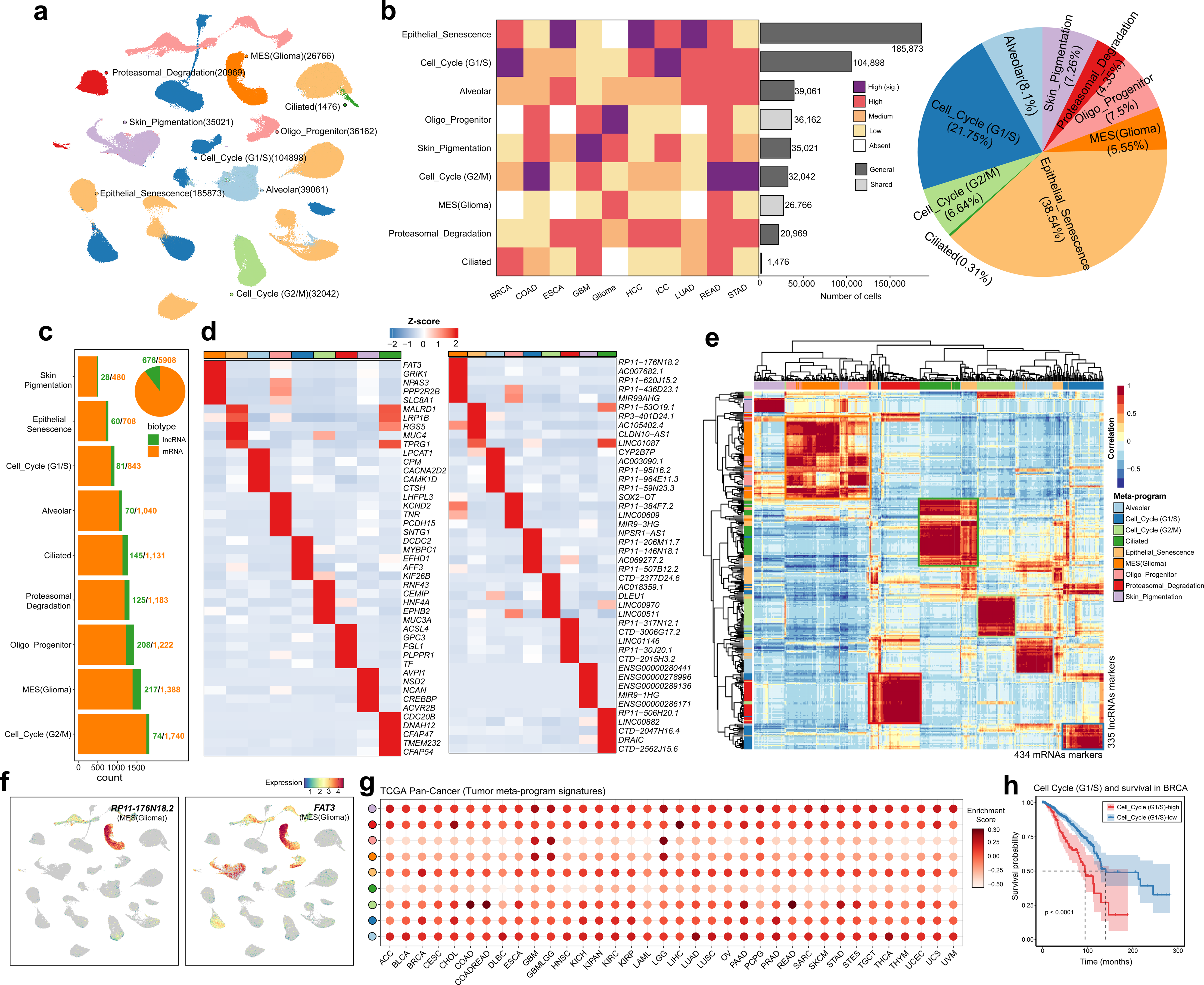
LncRNA-associated malignant meta-programs. **a** UMAP plot showing the distribution of MPs in each malignant cell. **b** Abundance of each MP (row) in each cancer type (column), defined as absent, low, medium, high or high and significant (left). Bar plot showing the number of cells in each MP (middle). MPs are further divided into two abundance categories by their abundance in each cancer type. Pie plot showing the proportions of each MP in all malignant cell (right). **c** Visualization of the number of lncRNA and mRNA markers in each MP (bar plot) and the total summary of markers (pie plot). **d** Heatmaps displaying top five mRNA (left) and lncRNA (right) markers for each MP based on their normalized expression profiles. **e** Heatmap of the similarity of lncRNA (rows) and mRNA (column) markers in each MP. The correlations were calculated by their expression profiles in each cluster. mRNA and lncRNA markers in each MP showed high similarity as boxed in the plot. **f** UMAP showing the expression patterns for selected lncRNA (*RP11-176N18.2*) and mRNA (*FAT3*) markers for MES (Glioma). The rest markers were visualized in Supplementary Fig. 3d-e. **g** Enrichment of MP signature in TCGA 36 primary cancers. Enrichment scores were shown both with color and size of points. Colored points indicate different MPs as shown in Figure 4A. **h** Kaplan-Meier plot comparing survival probability (Y-axis) across time (X-axis) between BRCA tumor in cell cycle (G1/S)-high and –low categories (colored), with p-value computed by log-ranked test. Error bands represent 95% confident intervals. The rest of the survival analyses were shown in Supplementary Fig. 4.

The MPs encompass crucial biological processes such as cell cycle (G1/S, G2/M) and epithelial senescence^5,8,9^. Notably, the MP associates with epithelial senescence exhibited a broader variability than anticipated, with widespread distribution observed in across BRCA, ESCA, HCC, ICC and LUAD (Fig. 4a). In order to gauge the prevalence of these robust MPs, we categories the frequency of each MP within each cancer type as absent, low, medium, high and significantly high (See Methods, Fig. 4b). Six MPs, including epithelial senescence, cell cycle (G1/S, G2/M), alveolar, skin pigmentation and proteasomal degradation, are consistently found at high or significantly high levels across multiple cancer types, thereby being classified as general MPs. In contrast, the remaining three MPs, detected at high levels in more than two cancer types, are denoted as shared MPs. The proportions of each MP range from 0.31% to 38.54%, highlighting the diverse landscape of MPs across different cancer types and underscoring the intricate interplay of biological processes contributing to cellular heterogeneity in cancer.

We further identify 6,908 mRNA and 676 lncRNA signatures in each MP (Fig. 4c, Supplementary Data 5). The number of mRNAs and lncRNAs associated with each MP range from 480 to 1,740 and 1,388, respectively. Notably, we observe that the abundance of mRNAs and lncRNAs in the MES (Glioma) and oligo progenitor clusters surpasses those in other MPs, potentially underpinning their heightened tissue specificity. Furthermore, we meticulously scrutinize the expression profiles of mRNAs and lncRNAs (Fig. 4d) and, remarkably, lncRNAs exhibit specificity scores on par with those of mRNAs. A subset of mRNAs and lncRNAs display pronounced MP specificity (Fig. 4f, Supplementary Fig. 3d-e). For instance, mRNAs such as *DCDC2*, *MYBPC1*, *EFHD1* and *AFF3* exhibit specific expression in the cell cycle (G1/S) cluster, and concomitantly, corresponding lncRNAs including *NPSR1-AS1*, *RP11-206M11.7*, *RP11-146N18.1*, *AC069277.2* and *RP11-507B12.2* are also markedly upregulated in the context of the cell cycle (G1/S). Collectively, our results suggest the pervasive cell type-specific expression patterns of lncRNAs across the entire context of malignant ITH and the role of lncRNAs in shaping the cellular landscape in cancer.

To establish the potential relationship between lncRNA signature and malignant programs, we conduct an extensive correlation analysis between a comprehensive set of lncRNAs (n = 676) and mRNAs (n = 6,908). Our investigation leads to the identification of 335 lncRNAs that demonstrated significant correlation (correlation coefficient > 0.7 and *P* value < 0.01) with 434 mRNAs (Fig. 4e). Moreover, our analysis reveals that the correlation between mRNAs and lncRNAs exhibited high similarity within each MP, further validating the possibility of utilizing these lncRNAs as promising candidate markers for malignant ITH programs. Moreover, we seek to ascertain whether the aforementioned MP enrichment patterns are recurrent across diverse primary cancers, which we accomplish by subjecting bulk sequencing data from the TCGA primary cancer database to pan-genome analysis. Our results reveal that the MP patterns are largely analogous across various primary cancers, as evidenced by a sizeable cohort of 12,193 samples (Fig. 4g). Subsequently, we endeavor to determine whether these MPs could be harnessed to prognosticate cancer survival. To this end, we conduct survival analyses of the MPs and found that all MPs are significantly associated with unfavorable clinical outcomes (Fig. 4h, Supplementary Fig. 4), underscoring the pivotal roles of lncRNAs in conferring malignant programs and directing them towards adverse clinical outcomes.

### Construction of lncRNA-associated networks in meta-programs

To gain a more nuanced understanding of the functional significances of lncRNA signatures in driving distinct tumor ITH programs, we seek to investigate the activity levels of transcription factors (TFs) and lncRNAs using SCENIC analysis^32^. Our analysis yields 468 significant regulons comprising a total of 19,568 genes across the nine MPs, with the number of genes varying between 98 and 326 in each MP (Fig. 5a, Supplementary Data 6). The results suggest the complex interplay between lncRNAs and TFs that govern cellular differentiation and malignant ITH in cancer.

**Fig. 5.**
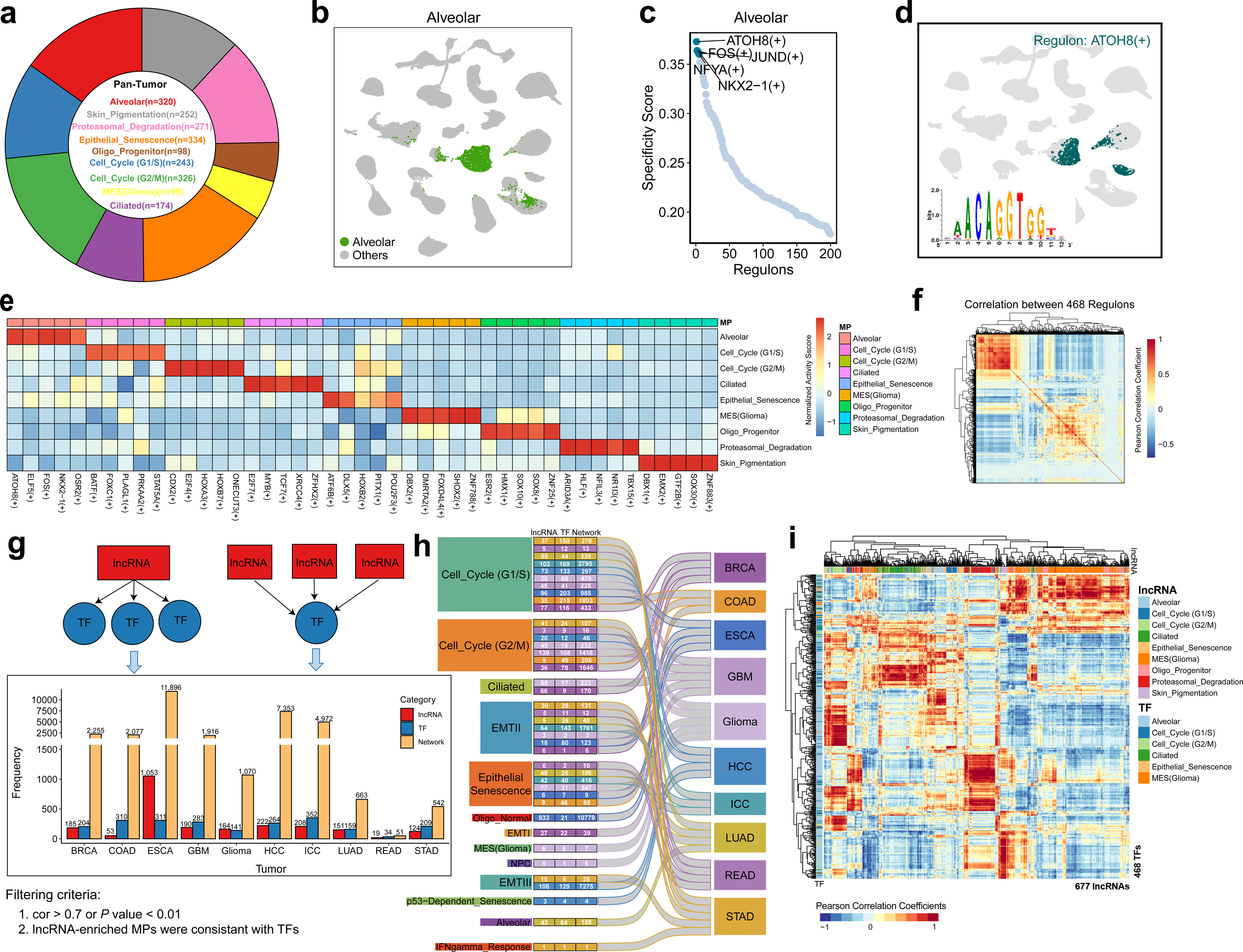
LncRNA-TF co-expression networks in tumor MPs. **a** Pie plot showing the number of TF regulators enriched in different tumor MP. **b** A highlighted MP (Alveolar) in the UMAP clusters. **c** Specificity score of regulons in Alveolar, with top five were labelled. **d** The UMAP showing the selected enriched *ATOHB* regulon. The rest of the top regulons enriched in each MP were shown in Supplementary Fig. 5. **e** Heatmap of top five regulons with the highest regulon activity in each MP. **f** Heatmap of the similarities of the 468 regulons. **g** The frequencies of lncRNAs, TFs and lncRNA-TFs networks in each tumor type. Two lncRNA-TF interaction modules were shown (top). **h** The numbers of lncRNAs, TFs and lncRNA-TF networks in each MP (left) and tumor type (right). **i** Heatmap of the correlation between significant lncRNA and TFs. Only *P* value < 0.01 or cor > 0.875 were retained for construction lncRNA-TF co-expression networks.

The above SCENIC analysis unveils some transcriptional regulons with remarkable specificity scores within distinct malignant ITH programs (Fig. 5b-d, Supplementary Fig. 5, Supplementary Data 6). Notably, the activity score-ranked regulons exhibit striking similarity to the specificity score-ranked regulons. Subsequently, we delve into the top five regulons in each MP (Fig. 5e). Consistent with the observed expression patterns, the alveolar programs demonstrate heightened activities of *ATOH8*, *NKX2-1*, *FOS*, *JUND* and *NFYA* regulons, while the *BATF* regulon is specifically enriched in the cell cycle (G1/S) program. Furthermore, the ciliated program displays distinct expression patterns of *E2F7*, *MYB*, *TCF7*, *XRCC4* and *ZFHX2* regulons, which have not been systematically investigated in epithelial cells previously. Additionally, we examine the transcriptional similarity of the regulons within each MP (Fig. 5f), revealing relatively low similarity between the regulons across different programs, underscoring their context-specific characteristics. To gain a deeper understanding of the functional roles of lncRNAs in transcriptional regulatory and the cooperative nature of different lncRNA-TF networks in cell fate specification, we employe a Connection Specificity Index (CSI) approach (See Methods) to quantify the functional similarities between lncRNA signatures and significant regulons. Following a rigorous filtering process, we obtain a set of significant lncRNAs, TFs and networks for each cancer types under investigation. For instance, in BRCA, we identify 185 significant lncRNAs, 204 TFs and 2,255 networks. To investigate the co-expression patterns between lncRNAs and TFs within their corresponding MPs, we perform correlation analysis on a joint dataset comprising 677 lncRNAs and 468 TFs. Notably, our analysis reveals high similarities in the co-expression profiles of these lncRNAs and TFs, supporting the notion of coordinated regulation within specific cellular contexts (Fig. 5i). In order to visualize the co-expression networks between lncRNAs and TFs, we filter and generate two types of co-expression networks by specific criteria of a lncRNA significantly co-expressing with more than 5 TFs or a TF significantly co-expressing with more than 10 lncRNAs (Supplementary Fig. 6). For example, within the cell cycle (G1/S) program, ten lncRNAs (*RP11-95P11.4*, *RP11-798K3.2*, *RP11-476K15.1*, *RP11-475O6.1*, *RP11-405A12.2*, *RP11-2337A12.1*, *CTD-2015H3.2*, *AP000472.2*, and *ADORA2A-AS1*) are found to act cooperatively in regulating a TF (*ZNF697*). Similarly, five TFs (*STAT5A*, *SP2*, *NR1I2*, *MSX2*, and *AHCTF1*) are regulated by a lncRNA (*RP4-798P15.3*). These results indicate the regulatory logic underlying the interactions between lncRNAs and TFs within the intricate gene regulatory networks governing diverse malignant cell activities.

## Discussion

In this study, we conduct a systematic analysis of random-primed snRNA-seq datasets, providing an unprecedented comprehensive atlas of lncRNAs within the context of tumor ITH programs in human cancers. Despite the numerous advantages of scRNA-seq technologies, their applicability in studying lncRNAs has been limited. Unlike many other scRNA-seq methods, the random-primed technologies enable the detection of total RNAs, making it particularly suitable for the analysis of lncRNAs^19,20^. Through this approach, we bridge the gap and generate novel insights into the involvement of lncRNAs in tumor heterogeneity and progression.

We develop scAnnotation, a novel method that combines dataset-based and marker-based algorithms to generate a robust reference atlas and cell taxonomies for single-cell technologies relying on random primers. Our approach allows for cell annotation using different references and specific tissue types, enabling the creation of a pan-cancer atlas covering 39 distinct major and rare cell types. Our analysis stratifies these cell types into metapopulations and provide a fine-grained map of TME cell states, i.e., a comprehensive view of cellular diversity within the TME. We expect that this approach will enable researchers to better understand the cellular diversity within complex biological systems, providing a foundation for the development of new therapeutic strategies for a wide range of disease.

Utilizing our atlas and pipeline, we have successfully generated a reference catalog containing well-defined cell states and their corresponding marker genes. The unprecedented scale of the datasets generated from the random-primed platform has enabled us to elucidate previously undescribed and overlooked cell markers, including a significant number of lncRNAs. Notably, our analysis has revealed novel new markers for specific cell clusters, which are not previously reported in the literature. To validate the robustness of these markers, we conduct comprehensive analyses using various commercial datasets, demonstrating their reliability and reproducibility across different tumor types and patients. Therefore, these markers hold promise for advancing our understanding of cellular heterogeneity in diverse biological contexts. Furthermore, we also observe the new specific expression of established known markers. For instance, *VWF*, traditionally known as endothelial cell marker^33^, is found to be specifically expressed in cluster 6 (VEC), highlighting the potential existence of subtype-specific expression patterns for canonical markers. Similarly, the master regulator of the mast cell lineage, *KIT*^34^, is specifically expressed in cluster 31 (mast cell). Our study not only expands the repertoire of known cell markers but also underscores the importance of rigorously validating these markers across diverse datasets and experimental settings. These results lay the groundwork and resource for future investigation aiming to unravel the functional significance of these markers in the context of complex biological systems and disease pathogenesis.

Numerous prior investigations have utilized low-throughput full-length single-cell RNA sequencing technologies to elucidate the roles of lncRNAs within various TME^11,12,35–37^. These studies have provided a comprehensive understanding of the complex landscape of the tumor-infiltrating environment. However, despite these advancements, a systematic overview of lncRNA-associated cell state determination and the identification of lncRNA markers for individual cell compositions across pan-cancer TME remains enigmatic. In this study, we have taken advantage of the random-primer approach by grouping lncRNAs based on their correlation with mRNA markers using hierarchical clustering, thereby expanding upon previous researches^12,35^. Notably, a prominent characteristic of lncRNAs is their tissue specificity, and our observations have revealed that the expression patterns of lncRNAs within each cluster exhibited high similarity with corresponding mRNAs. This underscores the highly cell– and tissue-specific nature of lncRNA expression. This comprehensive catalog of lncRNAs and their associations with distinct cell types are expected to serve as a valuable resource for future investigations into the roles of lncRNAs in pan-cancer TME cell state determination, as well as the cell type-specific functions of individual lncRNAs.

In order to gain a deeper understanding of lncRNA regulation in pan-cancer TME, we develop a novel tool called LncPairs, which allows us to identify cell type– or tissue-specific lncRNA-mRNA pairs at the single-cell level. Previous studies have highlighted the importance of cell-cell communication and crosstalk in multicellular organisms, such as human and mice^39^. Inspired by the principle of cell-cell crosstalk, we calculate the relationship between lncRNAs and mRNAs by employing cosine similarity analysis. In addition to confirming previously reported cell type-related lncRNA regulations, we also discover novel lncRNA-mRNA pairs. These findings provide valuable insights into the role of lncRNAs and lncRNA-mRNA pairs in determining cell states within the TME. Our study highlights the potential of lncRNAs as key regulators in cellular processes and emphasizes the importance of investigating lncRNA-mediated mechanisms for a comprehensive understanding of TME.

Previous studies have demonstrated that malignant cells cannot be clearly classified into distinct cell types, but rather exhibit diverse ITH programs^6–9^. Our study supports this notion and reveals that malignant cells exhibit expression patterns characterized by the activation of combinations of modules, indicating a continuum of variation rather than discrete clusters. However, we also observe instances where distinct states emerge when certain gene modules are mutually exclusive with others, exemplified by the ciliated, oligo progenitor, and MES (Glioma) programs, with the former being consistent with previous report^39^. By employing comprehensive analysis, we identify nine malignant programs, among which six display striking similarities to the heterogeneity programs observed within tumors. Our study highlights the significance of those programs and conservation even in the absence of a native microenvironment. The recurrence of these programs has been proposed as an indicator of their relevance^8,9,39^. Building upon this concept, we speculate that the recurrent programs identified in our study may exert important influences on cancer phenotypes such as tumor progression and metastasis. The intricate characterization of these malignant programs and the insights gained into their potential implications in cancer biology enhance our understanding of tumor heterogeneity.

Malignant cell states have been extensively studied, but the systematic identification of lncRNAs associated with these states remain incomplete. In this study, we identify a total of 676 lncRNA signature genes that serve as valuable resource for further investigation into their functions in various malignant cell states. Since lncRNAs generally interact with protein-coding genes and highly correlated genes tend to have similar functions, co-expression analysis of these lncRNAs with protein-coding genes can predict their potential functions or roles as markers. Furthermore, our single-nucleus transcriptome analyses demonstrate that individual malignant programs in pan-cancer environment can exhibit high expression of lncRNAs, consistent with above observations in different cell types. We also observe preferential dilution of certain lncRNAs in specific malignant programs, suggesting that state-specific expression of lncRNAs contribute to cellular identity. These analyses provide a resource of lncRNA signatures associated with malignant cell states and sheds light on their potential functions and markers.

The precise regulation of cellular differentiation is crucial for the establishment of diverse malignant programs. This process involves the differential expression genes, which is typically orchestrated by multiple TFs working in a coordinated manner^40,41^. By leveraging the power of an integrated single-nucleus transcriptome atlas encompassing various malignant cell states, we are able to evaluate the roles of key TFs in driving cell fate specification during malignant program development. The resulting program-specific gene GRNs provide valuable insights and generate hypotheses that can be further explored to elucidate the specific mechanisms underlying the formation of diverse malignant programs. For instance, *ATOH8*, *NKX2-1*, *FOS*, *JUND*, and *NFYA* have been well-established as regulators controlling the development and maintenance of alveolar epithelial cell identities^41–43^. Our analysis reveals the specific enrichment of these regulons within the alveolar program, strengthening their significance in driving the characteristics of this particular cell state. Interestingly, we observe that *BATF*, previously identified as an essential differentiation checkpoint in CD8^+^ T cells^44^, exhibits specific enrichment within the cell cycle (G1/S) program. This finding highlights the potential involvement of *BATF* in regulating cell cycle progression within malignant cells.

Given the growing recognition of lncRNAs in regulating cell type-specific gene expression, it is pertinent to systematically construct lncRNA-associated GRNs for lncRNAs. Our study identifies numerous transcriptional regulons and lncRNA-TF pairs that are likely to play distinct roles in driving the formation of diverse malignant programs. However, the functions of these TFs and lncRNAs, as well as their regulatory relationships with potential target genes, require extensive validation. One major challenge in this regard is the lack of appropriate experimental systems that can recapitulate the complexity of malignant programs. To address this issue, future studies need to leverage in vivo functional perturbation approaches to dissect the core transcriptional regulators and lncRNA-TF regulatory relationships in transgenic animals. Such studies will enable the characterization of transcriptional and phenotypic changes at high spatiotemporal resolution, providing valuable insights into the complex regulatory networks underlying malignant ITH programs. Furthermore, the use of advanced imaging techniques, such as intravital microscopy, will reveal the dynamics of gene expression and cellular behavior within the TME, shedding light on how different cell states interact and coexist during malignant program development.

In conclusion, this study represents a significant advancement in our understanding of the roles of lncRNAs in single-nucleus pan-cancer TME and malignant programs. We have generated a comprehensive catalog of lncRNAs and their functional repertoires in various malignant cell states, complemented with useful tools/pipelines for further analysis. Our results provide compelling evidence that a multitude of novel lncRNAs are present within the pan-cancer tumor cell and are differentially involved in regulating the expression of numerous genes in distinct malignant programs. Our study provides valuable resource for future investigations into the functional roles of lncRNAs in ITH-based research.

## Methods

### Data collection and processing

The processed snRNA-seq datasets in the form of raw counts are acquired from our previous studies^20,21^. In total, 49 datasets comprising 49 samples are obtained encompassing ten distinct cancer types (Fig. 1a). Upon acquisition, each dataset’ raw unique identifier (UMI) count matrices in .h5ad format are initially converted into Seurat^45^ object in .rds format, independently. This conversion facilitates downstream analysis within the R environment.

To ensure the accuracy and reliability of our analysis, we subject the original count matrices to rigorous quality control (QC) process based on three metrics. Firstly, we filter cells with low quality based on their total UMI counts, number of detected genes, and mitochondrial RNA percentage. Specifically, cells with less than 500 UMIs and fewer than 200 detected genes are excluded, as well as those with mitochondrial gene percentages exceeding 30%. Additionally, to eliminate any potential doublets, we remove cells with an unusually high number of UMIs (> 30,000) and detected genes (> 5,000). Following this QC process, a total of 801,879 clean cells remains for downstream analysis.

### Integration of multiple snRNA-seq datasets

To address the challenge of batch effects within single-nucleus RNA sequencing datasets of ten different cancer types, we employe a two-round approach using the Harmony algorithm^46^, which is specifically designed for integrating large-scale single-cell datasets. In the first round of Harmony integration, we focus on removing batch effects associated with two different snRNA-seq protocols: snHH-seq and snRandom-seq. By applying the algorithm, we effectively mitigate the batch effects arising from these two distinct experimental procedures. Subsequently, we perform a second round of integration to address the batch effects originating from the diverse cancer types included in our study. The application of Harmony in this round facilitates the removal of batch effects associated with the different cancer types, enabling a more accurate representation of the underlying biological variation. The outputs obtained from the integration and batch effect correction steps serve for downstream analysis, including dimensional reduction and cell type annotation.

### Dimension reduction and unsupervised clustering

Subsequently, the integrated datasets undergo comprehensive processing for dimensional reduction and unsupervised clustering according to the workflow implemented in Seurat^45^. Initially, raw counts are subjected to normalization using the *NormalizeData* function, with the parameter “*normalization.method = ‘LogNormalize’*, *scale.factor = 10000*”. Following normalization, a selection of 2,000 highly-variable genes is identified for downstream analysis through the utilization of the *FindVariableFeatures* function, employing the parameter “*selection.method = ‘vst’, nfeatures = 2000*”. To mitigate the effects of total count per cell and the percentage of mitochondrial gene count, the *ScaleData* function is applied with the parameter “*vars.to.regress = c(’nCount_RNA’, ‘percent.mito’)*” to regress out these confounding factors. Subsequently, a principal component analysis (PCA) matrix comprising 50 components is computed to unveil the primary axes of variation and denoise the data, achieved through the implementation of *RunPCA* function with parameters “*features = VariableFeatures(object), npcs = 50*”. The optimal number of principal components is determined based on three metrics: 1) cumulative contribution of principal components greater than 90%; 2) individual principal components contributing less than 5% to variance; and 3) a difference of less than 0.1% between two successive principal components. Dimensional reduction is then executed by employing the UMAP algorithm, offering a two-dimensional embedding for data visualization. Subsequent cluster analysis is conducted using the Leiden algorithm, a method specifically designed for identifying communities or clusters within a network by optimizing a quality function that measures the modularity of the network.

### Cell type annotation

To expand the landscape of TME beyond the existing pan-cancer datasets using single-cell and sc/snRNA-seq, we develop an unsupervised cell type annotation tool called scAnnotation. This tool employs two distinct strategies for cell type identification, including a dataset-based method and a marker-based method (Supplementary Fig. 1a). For the dataset-based approach, we utilize the SingleR^47^ algorithm. However, the in-built reference datasets provided by SingleR may not fully meet the requirements for accurate cell type annotation. To address this limitation, we create well-annotated reference datasets which are specifically designed to enhance the accuracy of cell type identification in scAnnotation. In addition to the dataset-based approach, we also implement a marker-based strategy by integrating markers from multiple sources, including ScType^48^, CellMarker^49^, and CellTaxonomy^50^. By combining markers from these diverse sources, we aim to increase the robustness and accuracy of our cell type identification. To ensure the validity of our cell type annotations, we manually curate the results generated by both author-based annotations and scAnnotation-defined annotations. This curation process allows us to identify potential errors or inconsistencies in our cell type assignments and rectify them accordingly.

### Identification of marker genes

In order to evaluate the effectiveness of our integrated single-nucleus transcriptomic map in identifying rare cell types and constructing a marker database for annotating datasets generated from random-primed snRNA-seq technologies, we implement a stringent workflow (Fig. 2a). To identify robust marker genes for each cell type, we employe a combination of differential expression analysis (DEA) and COSG^51^. For DEA, we utilize the FindMarkers function in Seurat^45^, with the following parameters: “*logfc.threshold = 0.25, min.pct = 0.1*”. Additionally, we apply COSG with the parameter “*mu = 1, n_gene_user = 500*”. To ensure the accuracy of identifying significant cluster markers, we perform DEA on the normalized RNA counts rather than the integrated data. To refine the cluster signatures further, we use both the Wilcoxon rank-sum test and COSG algorithm methods and retains only the genes that are identified by both approaches. These markers are then categorized into mRNA and lncRNA categories. For the identification of mRNA markers, we employe a multi-step approach. Firstly, we classify the cell types into tissue-shared and tissue-specific categories. Next, we perform DEA separately for each cell type within each category. Subsequently, we narrow down the number of markers through overlapping analysis at both the tissue and sample levels. Furthermore, we conduct functional enrichment analysis to further refine the set of marker genes. Only genes enriched in corresponding cell type terms (using GSEA, C8) are retained. Finally, we compare these markers with publicly available well-known marker databases such as CellMarker^49^, HCL^52^, PanglaoDB^53^, among others. To identify lncRNA markers, we perform correlation analysis between the curated mRNA markers and the previously obtained lncRNA markers. Only lncRNAs that exhibited high correlation (correlation > 0.875 or *P* value < 0.01; within the same cluster) with the mRNA markers are considered as lncRNA markers.

### Calculation of cell-lineage specificity

Cell lineage specificity (CSI) score for each gene was calculated as follows: 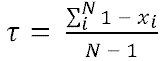, where *N* refers to the number of cell clusters as x_i_ denotes the expression level in cluster *i*, and is normalized by the maximal expression value across *N* clusters. Above a given threshold, within a cluster compared to outside a cluster. To generate the set of mRNAs and lncRNAs specific to each cell type, we first assign each gene to a cell type by determining which cell type yielded the maximum odds specificity score for that gene. We use each gene’s own 75%ile for expression thresholds. Then we calculate expression enrichment scores for each gene in each cell type by: 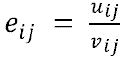 is the mean expression level of gene *i* within cluster *j*, and *v_i,j_* is the mean expression level of gene i outside of cluster *j*. Genes are ranked within their cell type cluster assignments using expression enrichment scores and the top 3 mRNAs and lncRNAs in each cell type are presented.

### Identification of lncRNA-mRNA pairs

Sometimes, only mRNA or lncRNA could not define cell state well, lncRNA-mRNA interaction pairs will be a new perspective to confer this work. We propose a lncRNA-mRNA interaction pairs computational workflow, LncPairs, which infers tissue– and cell-specific lncRNA-mRNA pairs at cell level. The main pipeline is shown in Fig. 3a, can be illustrated as follows: 1) Top 2,000 HVGs (highly variable genes)-based single-cell expression matrix is used to construct gene x cluster expression matrix. The expression of each gene is averaged based on clusters. 2) Splitting the gene x cluster matrix into mRNA x cluster and lncRNA x cluster matrices. 3) Calculating the correlation between the mRNA x cluster and lncRNA x cluster matrix. Only lncRNA-mRNA pairs with PCC (Pearson Correlation Coefficient) > 0.85 are remained for further investigation. 4) Constructing pair x cluster matrix by scoring the remained lncRNA-mRNA pairs, the scoring function is inspired by Armingol et al^38^. 5) Cosine similarity method is to identify cluster-specific lncRNA-mRNA pairs. 6) Filtering out pairs of similarity score less than 0.95.

### Functional enrichment analysis

To infer the biological functions of the markers, we develop a geneset-based functional enrichment method, called scEnrichment. In this method, we primarily use signatures from MsigDB, including the following collections of gene sets: Gene Ontology (BP, CC, and MF), Hallmark and KEGG. Only signatures with *P* value < 0.05 are considered significantly enriched.

### MP annotation

As tumor cell are highly heterogeneous, we next cluster malignant cell extracted from tumor environment based on robust NMF (Nonnegative Matrix Factorization) programs and Jaccard similarity. NMF algorithm is widely applied in single cell tumor research^6,8,9^. We carry out NMF implemented in RcppML package (https://github.com/zdebruine/RcppML) for each sample separately and integrate samples, to generate programs that capture the heterogeneity within each sample and tumor. Negative values in each centered expression matrix are set to zero. As application of NMF requires a ‘K’ parameter that influences the results, we run NMF using different values (K = 10, 11, 12, to 20) for each tumor dataset. The most suitable K is determined as the number of generated MPs was equal to the K.

We next cluster the robust NMF programs according to Jaccard similarity. Briefly, each NMF program is compared to reference MPs summarized from public researches^8,39^ to assess the degree of gene overlap between programs. The NMF program is defined by the following criteria: the overlap instances of at least 5 genes; the program with the maximal number of considerable overlaps.

### MP abundance estimation

We first calculate, for each MP, the observed proportions of MP programs in each cancer type. The abundance is then defined by quantiles. The abundance classification in Fig. 4c is defined as follows: High (significant): 100% MP programs in that cancer type; High: 75%-100% MP programs in that cancer type; Medium: 50%-75% MP programs in that cancer type; Low: 25%-50% MP programs in that cancer type and Absent: 0%-25% MP programs in that cancer type.

### Inference of MP regulators

SCENIC^32^ infers transcriptional regulators, which consist of a TF and its direct target genes, by incorporating both the co-expression patterns and the enrichment of TF binding motifs in regions upstream of transcription start sites (TSSs). Activated regulons in malignant cell subsets are analyzed by SCENIC with raw count matrix as input. The integrated malignant cell labelled with MPs is used as input for the python implementation of the SCENIC algorithm (pyscenic). Firstly, genes are filtered to have at least 0.05 counts per cell on average and to be detected in at least 5% of the cells, and the transcription factor-binding databases used are 500bp-upstream and transcription start site-centered-10kb. The gene-gene co-expression relationships between transcription factors (TF) and their potential targets are inferred using the *grn* function with the gene regulatory network reconstruction algorithm *GRNBoost2* and the regulon are identified by RcisTarget. Next, the regulon activity for each MP is scored by AUCell function. Briefly, GRN (gene regulatory network) contains three steps: 1) identify co-expression modules between TF and the potential targets; 2) calculate Regulon Activity Score (RAS) for each cluster.

To connect regulons with MPs, we use Jensen-Shannon Divergence (JSD) function implemented in philentropy package (https://github.com/drostlab/philentropy) to identify MP-specific regulons with AUC score matrices, and the corresponding binding motifs for regulons were obtained from JASPAR database. Binary RAS for regulons is projected to the regulon activity UMAP projection. We further calculate similarity of regulons to identify regulon modules that consist of multiple functionality related regulons. For each pair-wise combination of regulons, we calculate the Connection Specificity Index (CSI)^54^ as a measure for evaluating the extent of functional association between two partners. Hierarchical clustering with Euclidean distance is then performed on the CSI matrix of regulons. We use CSI > 0.7^55^ as a cutoff to identify regulon modules.

## Data availability

This work relies on curation and integrative analysis of published studies and do not involve generation of new primary data. The integrated UMI matrices and metadata are available at https://zenodo.org/records/10024207.

## Code availability

The code for cell type annotation is available at https://github.com/EternalVoice/scAnnotation. Code for inferring lncRNA-mRNA pairs is available at https://github.com/EternalVoice/LncPairs. Code for performing single cell enrichment analysis is available at https://github.com/EternalVoice/scEnrichment.

## Acknowledgments

We thank the technical support by the Core Facilities of Liangzhu Laboratory and the Core Facilities of Zhejiang University School of Medicine. Thanks for M20 Genomics bioinformatics team for the valuable discussion.

## Author contribution

Y.W., L.F. and T.F. conceived the study and designed the project. H.C. and Z.X. provided the sequencing datasets. S.N. provided technique support. Y.W. and L.F. supervised this project. T.F. performed data analysis and wrote the manuscript and all authors have revised and approved the final manuscript.

## Supplementary Data

**Supplementary Fig. 1. Identification and visualization of markers. a** Illustration of the bioinformatics pipeline for integrative analysis of single cell datasets. UMAP showing 801,879 single cells from pan-cancer samples grouped by cancers (**b**) and patients (**c**). **d** Pipeline to perform cell type annotation. **e** Pie plot showing the generated marker lists can be annotated by public marker databases. **f** UMAP visualizing of markers in each cell types, corresponding to Figure 1E.

**Supplementary Fig. 2. Visualization of lncRNA markers of each cell type. a** Heatmap showing the correlation of lncRNA (row) and mRNA (column) markers. The correlations were calculated by their expression profiles in each cluster. mRNA and lncRNA markers in each cluster showed high similarity. **b** Heatmaps displaying top 3 lncRNA markers for each cell type based on their cell type specificity scores. The most specific lncRNAs were labelled in red. **c** UMAP showing the expression profiles of lncRNA markers for each cell type. Correspond to Fig. 2d.

**Supplementary Fig. 3. Visualization of malignant MPs. a** Schema for defining and functionally annotating malignant MPs. **b** Jaccard similarity indices for comparisons among NMF programs based on their marker genes. **c** UMAP plot showing exemplary cancer types for the three MP defined by the presence and number of discrete subpopulations that were identified using NMF. UMAP showing expression patterns of mRNA (**d**) and lncRNA (**e**) markers of each MP, corresponding to Fig. 4f.

**Supplementary Fig. 4. TCGA survival analysis of each MP.** Clinical outcomes of each MP by TCGA pan-cancer datasets, corresponding to Fig. 4h.

**Supplementary Fig. 5. Top regulons that regulate transcriptional activity of MPs. a** UMAP showing distribution patterns of each malignant MP, correlating with Figure 5B. **b** Dot plot showing specificity score of regulons with top 5 are labelled, corresponding to Fig. 5c. **c** UMAP showing top regulons enriched in corresponding MP with motifs are labelled, correlating to Fig. 5d.

**Supplementary Fig. 6. LncRNA-TF regulatory networks. a** TF-oriented lncRNA co-expression networks of each MP. **b** lncRNA-oriented TF co-expression networks of each MP. For visualization of these networks, we applied the criteria: 1) one lncRNA significantly co-expressed with more than 5 TFs; 2) one TF significantly co-expressed with more than 10 lncRNAs.

## Reference

1. Marusyk, A., Almendro, V. & Polyak, K. Intra-tumour heterogeneity: a looking glass for cancer? Nat. Rev. Cancer 12, 323–334 (2012).

2. Vitale, I., Shema, E., Loi, S. & Galluzzi, L. Intratumoral heterogeneity in cancer progression and response to immunotherapy. Nat. Med. 27, 212–224 (2021).

3. Tirosh, I. et al. Dissecting the multicellular ecosystem of metastatic melanoma by single-cell RNA-seq. Science 352, 189–196 (2016).

4. Rambow, F. et al. Toward Minimal Residual Disease-Directed Therapy in Melanoma. Cell 174, 843–855 (2018).

5. Neftel, C. et al. An integrative model of cellular states, plasticity, and genetics for glioblastoma. Cell 178, 835–849 (2019).

6. Puram, S. et al. Single-cell transcriptomic analysis of primary and metastatic tumor ecosystems in head and neck cancer. Cell 171, 1611–1624 (2017).

7. Moncada, R. et al. Integrating microarray-based spatial transcriptomics and single-cell RNA-seq reveals tissue architecture in pancreatic ductal adenocarcinoma. Nat. Biotechnol. 38, 333–342 (2020).

8. Gavish, A. et al. Hallmarks of transcriptional intratumour heterogeneity across a thousand tumours. Nature 618, 598–606 (2023).

9. Kinker, G.S. et al. Pan-cancer single-cell RNA-seq identifies recurring programs of cellular heterogeneity. Nat. Genet. 52, 1208–1218 (2020).

10. Xu, J. et al. A comprehensive overview of lncRNA annotation resources. Brief Bioinform. 18, 236–249 (2017).

11. Hon, C.C. et al. An atlas of human long non-coding RNAs with accurate 5’ ends. Nature 543, 199–204 (2017).

12. Zhou, J. et al. Combined Single-Cell Profiling of lncRNAs and Functional Screening Reveals that H19 Is Pivotal for Embryonic Hematopoietic Stem Cell Development. Cell Stem Cell 24, 285–298 (2019).

13. Catellanos-Rubio, A. et al. A long noncoding RNA associated with susceptibility to celiac disease. Science 352, 91–95 (2016).

14. Zhao, H., Zhang, X., Frazão, J.B., Condino-Neto, A. & Newburger, P.E. HOX antisense lincRNA HOXA-AS2 is an apoptosis repressor in all trans retinoic acid treated NB4 promyelocytic leukemia cells. J. Cell Biochem. 114, 2375–2383 (2013).

15. Park, E.G., Pyo, S.J., Cui, Y., Yoon, S.H. & Nam, J.W. Tumor immune microenvironment lncRNAs. Brief Bioinform. 23, bbab504 (2022).

16. Zheng, G.X.Y. et al. Massively parallel digital transcriptional profiling of single cells. Nat. Commun. 8, 14049 (2017).

17. Salmen, et al. High-throughput total RNA sequencing in single cells using VASA-seq. Nat. Biotechnol. 40, 1780–1793 (2022).

18. Sheng, et al. Effective detection of variation in single-cell transcriptomes using MATQ-seq. Nat. Methods 14, 267–270 (2017).

19. Xu, Z. et al. High-throughput single nucleus total RNA sequencing of formalin-fixed paraffin-embedded tissues by snRandom-seq. Nat. Commun. 14, 2734 (2023).

20. Chen H, et al. Pan-Cancer Single-Nucleus Total RNA Sequencing Using snHH-seq. Adv. Sci. 2023, 2304755. doi: 10.1002/advs.202304755 (2023).

21. Xu Z. et al. An automated archival single-nucleus total RNA sequencing platform mapping integrative and retrospective cell atlas of glioma. bioRxiv doi: 10.1101/2023.11.16.567325 (2023).

22. Lin, W. et al. Single-cell transcriptome analysis of tumor and stromal compartments of pancreatic ductal adenocarcinoma primary tumors and metastatic lesions. Genome Med. 12, 80 (2020).

23. Wu, F. et al. Single-cell profiling of tumor heterogeneity and the microenvironment in advanced non-small cell lung cancer. Nat. Commun. 12, 2540 (2021).

24. Muhl, L. et al. Single-cell analysis uncovers fibroblast heterogeneity and criteria for fibroblast and mural cell identification and discrimination. Nat. Commun. 11, 3953 (2020).

25. Su, X. et al. Comprehensive analysis of long non-coding RNAs in human breast cancer clinical subtypes. Oncotarget 5, 9864–9876 (2014).

26. Mamivand, A. et al. Data mining of bulk and single-cell RNA sequencing introduces OBI1-AS1 as an astrocyte marker with possible role in glioma recurrence and progression. Clin. Epigenetics 14, 35 (2022).

27. Min, S.Y. et al. Multiple human adipocyte subtypes and mechanisms of their development. bioRxiv 10.1101/537464 (2019).

28. Boos, F. et al. The endothelial-enriched lncRNA LINC00607 mediates angiogenic function. Basic Res. Cardiol. 118, 5 (2023).

29. Zeng, W. et al. CCL18 signaling from tumor-associated macrophages activates fibroblast to adopt a chemoresistance-inducing phenotype. Oncogene 42, 224–237 (2023).

30. Li, J. et al. VSIG4 inhibits proinflammatory macrophage activation by reprogramming mitochondrial pyruvate metabolism. Nat. Commun. 8, 1322 (2017).

31. Wiggins, K.A. et al. IL-1α cleavage by inflammatory caspases of the noncanonical inflammasome controls the senescence-associated secretory phenotype. Aging Cell 18, e12946 (2019).

32. Aibar, S. et al. SCENIC: single-cell regulatory network inference and clustering. Nat. Methods 14, 1083–1086 (2017).

33. Nakhaei-Nejad, M., Farhan, M., Mojiri, A., Jabbari, H., Murray, A.G. & Jahroudi, N. Regulation of von Willebrand Factor Gene in Endothelial Cells That Are Programmed to Pluripotency and Differentiated Back to Endothelial Cells. Stem Cells 37, 542–554 (2019).

34. Tasi, M., Valent, P. & Galli, S.J. KIT as a master regulator of the mast cell lineage. J. Allergy Clin. Immunol. 149, 1845–1854 (2022).

35. Bjørklund, S.S. et al. Subtype and cell type specific expression of lncRNAs provide insight into breast cancer. *Commun*. Biol. 5, 834 (2022).

36. Aghagolzadeh, P. et al. Assessment of the Cardiac Noncoding Transcriptome by Single-Cell RNA Sequencing Identifies FIXER, a Conserved Profibrogenic Long Noncoding RNA. Circulation 148, 778–797 (2023).

37. Zhou, M. et al. The transcriptional landscape and diagnostic potential of long non-coding RNAs in esophageal squamous cell carcinoma. Nat. Commun. 14, 3799 (2023).

38. Armingol, E., Officer, A., Harismendy, O. & Lewis, N.E. Deciphering cell-cell interactions and communication from gene expression. Nat. Rev. Genet. 22, 71–88 (2021).

39. Barkley, D. et al. Cancer cell states recur across tumor types and form specific interactions with the tumor microenvironment. Nat. Genet. 54, 1192–1201 (2022).

40. Reiter, F., Wienerroither, S. & Stark, A. Combinatorial function of transcription factors and cofactors. Curr. Opin. Genet. Dev. 43, 73–81 (2017).

41. Zhou, B. et al. Comprehensive epigenomic profiling of human alveolar epithelial differentiation identifies key epigenetic states and transcription factor co-regulatory networks for maintenance of distal lung identity. BMC Genomics 22, 906 (2021).

42. Watanabe, N. et al. Anomalous Epithelial Variations and Ectopic Inflammatory Response in Chronic Obstructive Pulmonary Disease. Am. J. Respir. Cell Mol. Biol. 67, 708–719 (2022).

43. Little, D.R. et al. Transcriptional control of lung alveolar type 1 cell development and maintenance by NK homeobox 2-1. Proc. Natl. Acad. Sci. USA 116, 20545–20555 (2019).

44. Kurachi, M. et al. The transcription factor BATF operates as an essential differentiation checkpoint in early effector CD8^+^ T cells. Nat. Immunol. 15, 373–383 (2014).

45. Hao, Y.H. et al. Integrated analysis of multimodal single-cell data. Cell 184, 3573–3587 (2021).

46. Korsunsky, I. et al. Fast, sensitive and accurate integration of single-cell data with Harmony. Nat. Methods 16, 1289–1296 (2019).

47. Aran, D. et al. Reference-based analysis of lung single-cell sequencing reveals a transitional profibrotic macrophage. Nat. Immunol. 20, 163–172 (2019).

48. Ianevski, A., Giri, A.K. & Aittokallio, T. Fully-automated and ultra-fast cell-type identification using specific marker combinations from single-cell transcriptomic data. Nat. Commun. 13, 1246(2022).

49. Zhang, X.X. et al. CellMarker: a manually curated resource of cell markers in human and mouse. Nucleic Acids Res. 47, D721–D728 (2019).

50. Jiang, S. et al., Cell Taxonomy: a curated repository of cell types with multifaceted characterization. Nucleic Acids Res. 51, D853–D860 (2023).

51. Dai, Min., Pei, X.B. & Wang, X.J. Accurate and fast cell marker gene identification with COSG. Brief Bioinform. 23, bbab579 (2022).

52. Han, X. et al. Construction of a human cell landscape at single-cell level. Nature 581, 303–309 (2020).

53. Franzén, O., Gan, L.M. & Björkegren, J.L.M. PanglaoDB: a web server for exploration of mouse and human single-cell RNA sequencing data. Database (Oxford) 2019, baz046 (20219).

54. Bass, J.I.F, Diallo, A., Nelson, J., Soto, J.M., Myers, C.L. & Walhout, A.J. Using networks to measure similarity between genes: association index selection. Nat. Methods 10, 1169–1176(2013).

55. Suo, S., Zhu, Q., Saadatpour, A., Fei, L., Guo, G. & Yuan, G.C. Revealing the Critical Regulators of Cell Identity in the Mouse Cell Atlas. Cell Rep. 25, 1436–1445 (2018).

