## Supplementary figures and images for "Landscape and functional repertoires of long noncoding RNAs in the pan-cancer tumor microenvironment using single-nucleus total RNA sequencing"

### Supplemental Figure 1

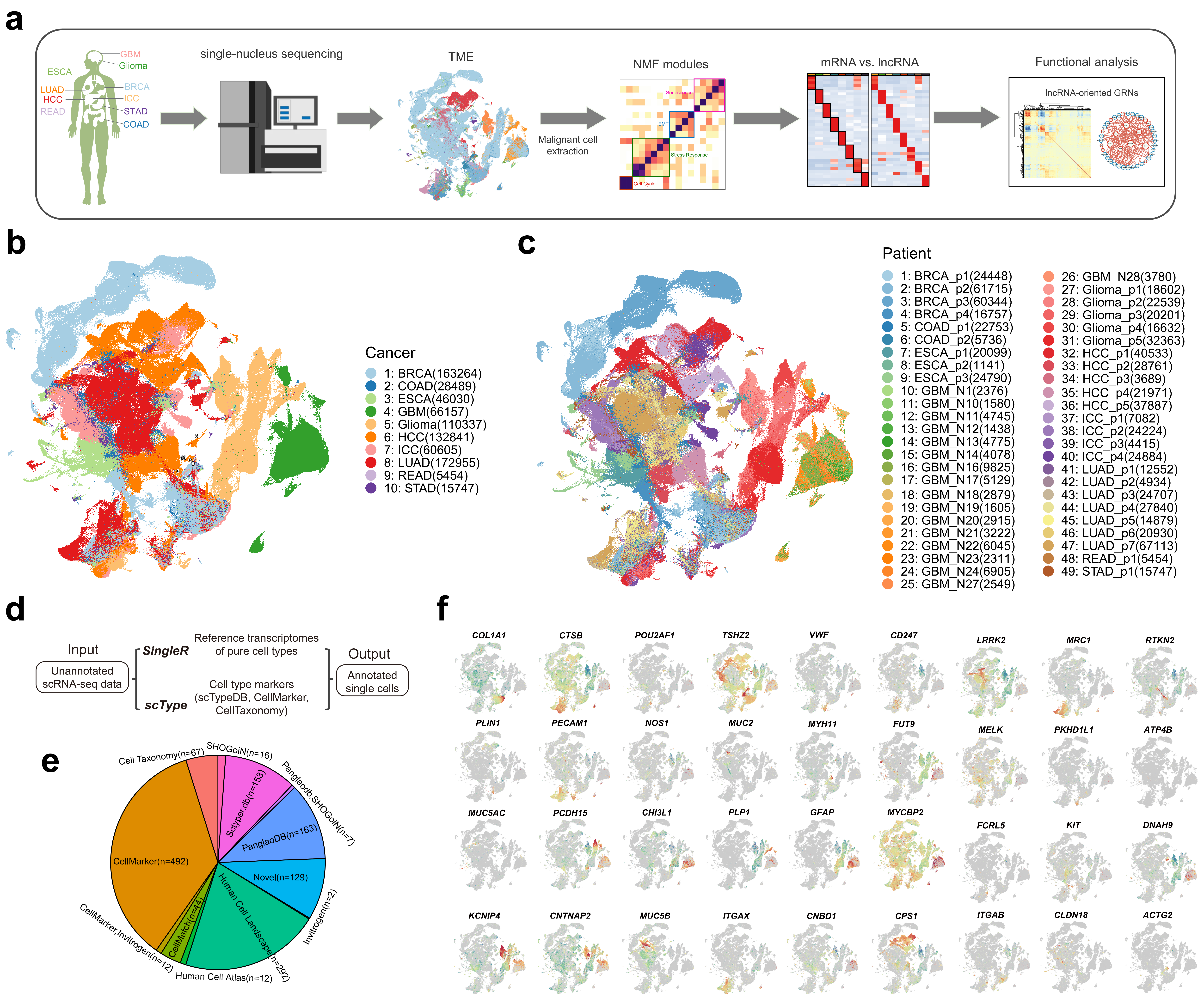

### Supplemental Figure 2

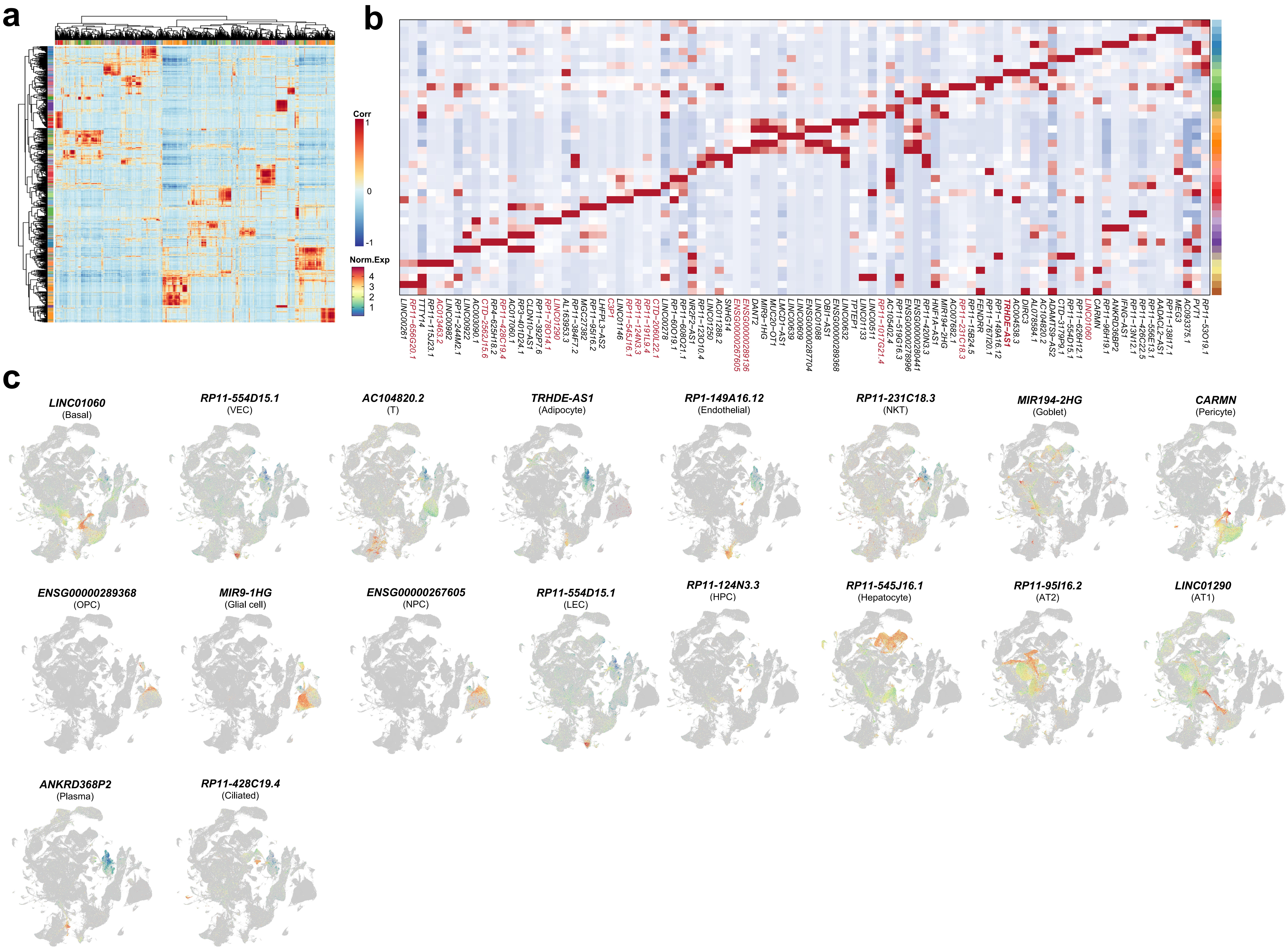

### Supplemental Figure 3

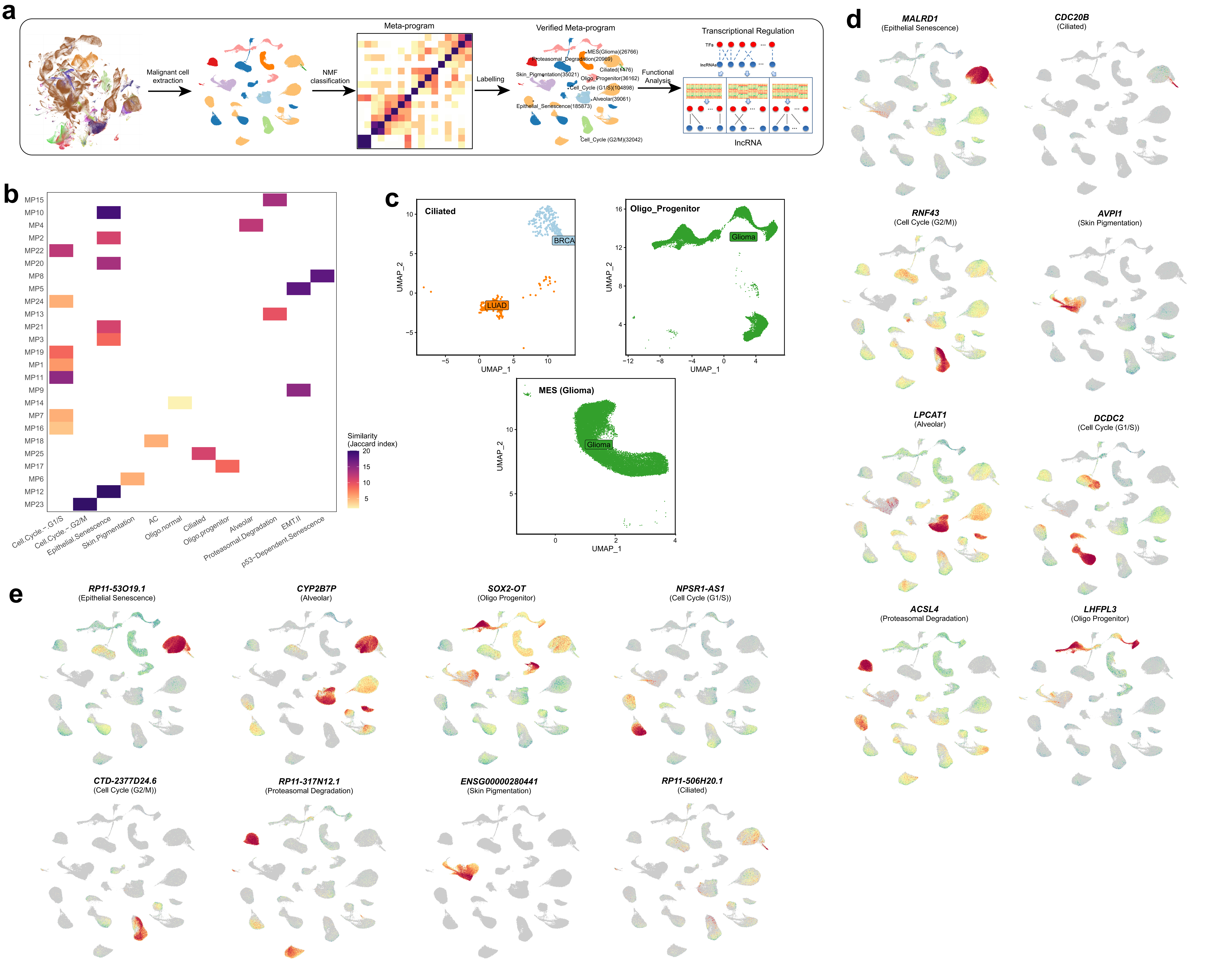

### Supplemental Figure 4

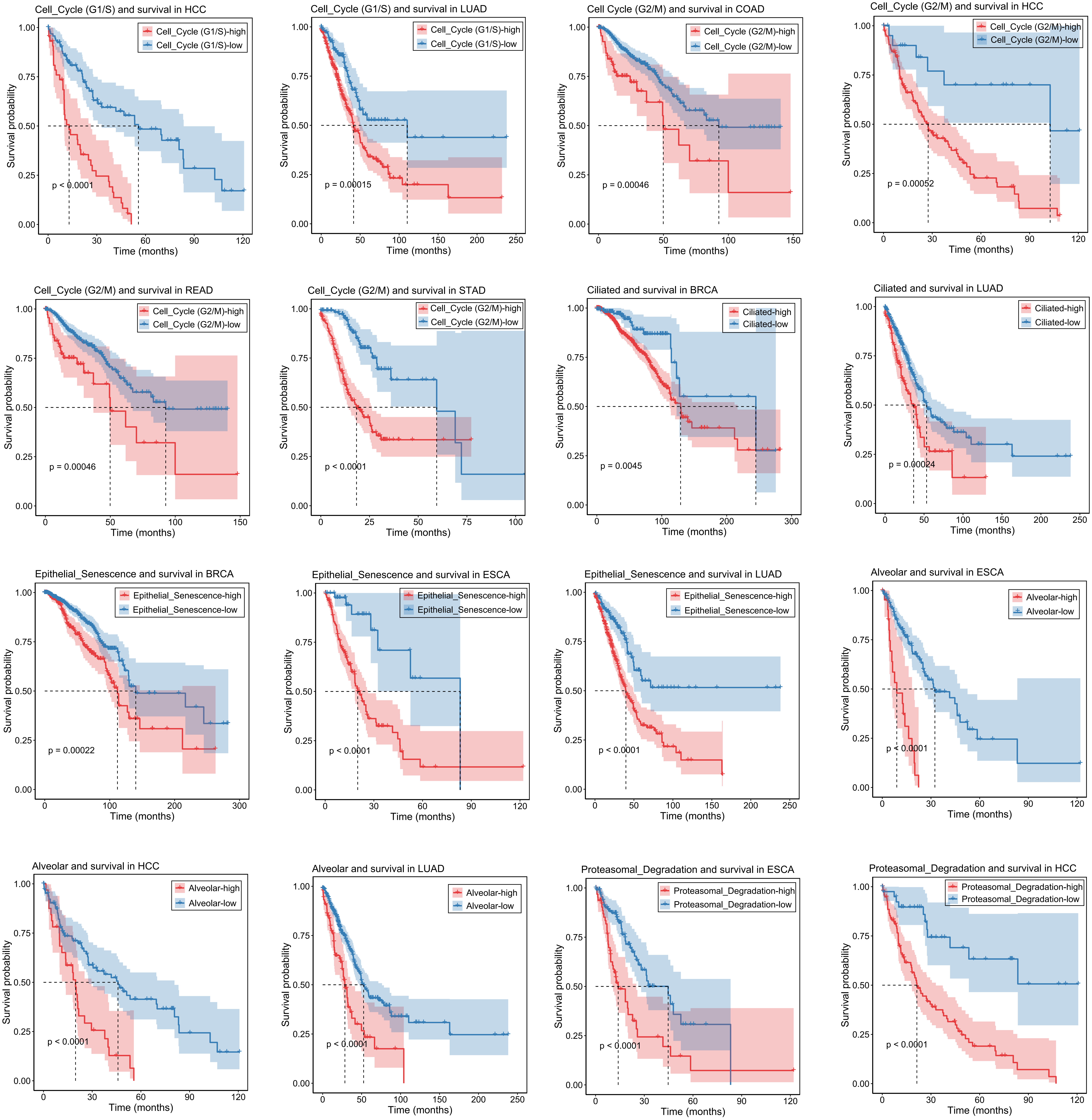

### Supplemental Figure 5

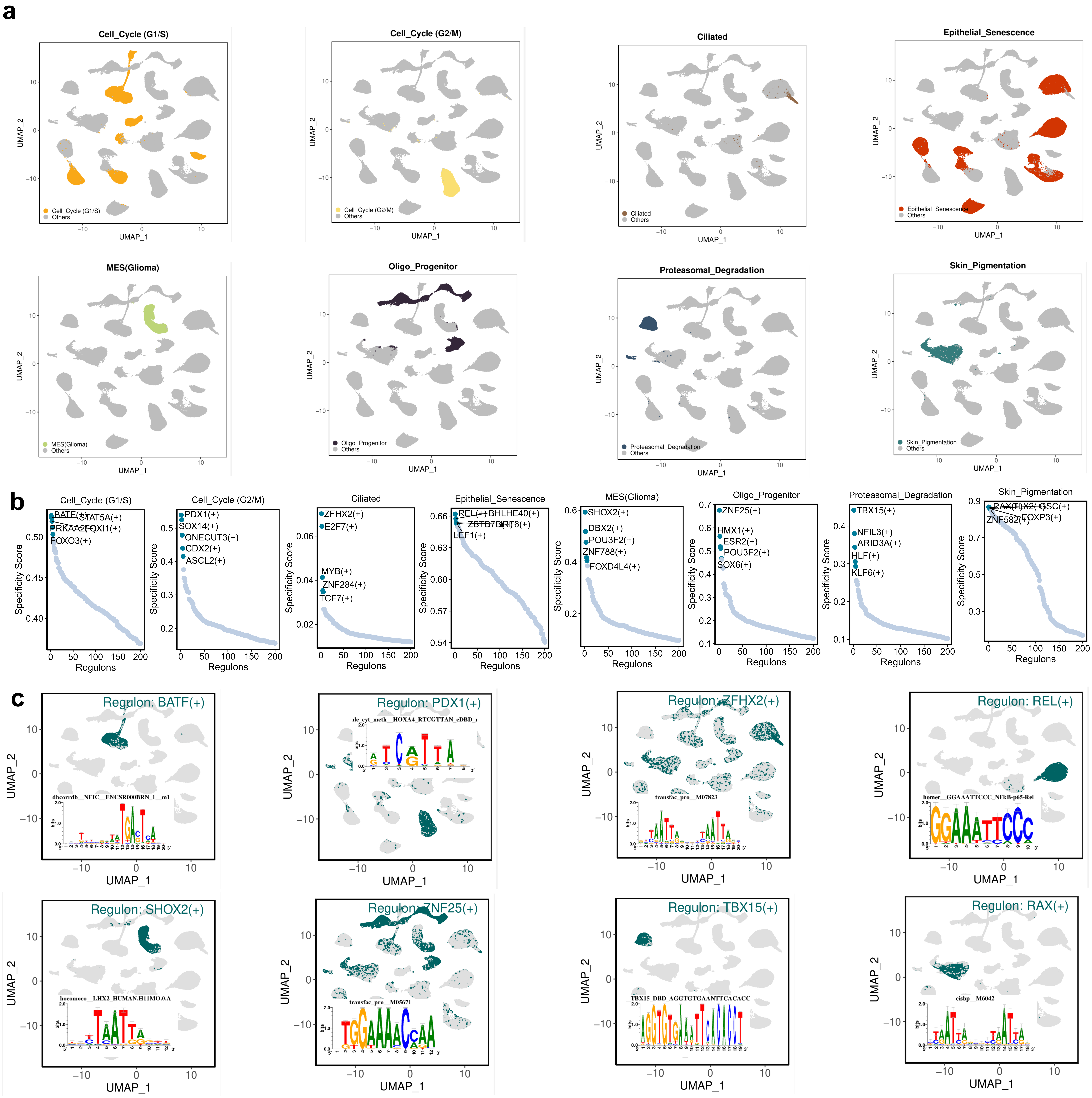

### Supplemental Figure 6

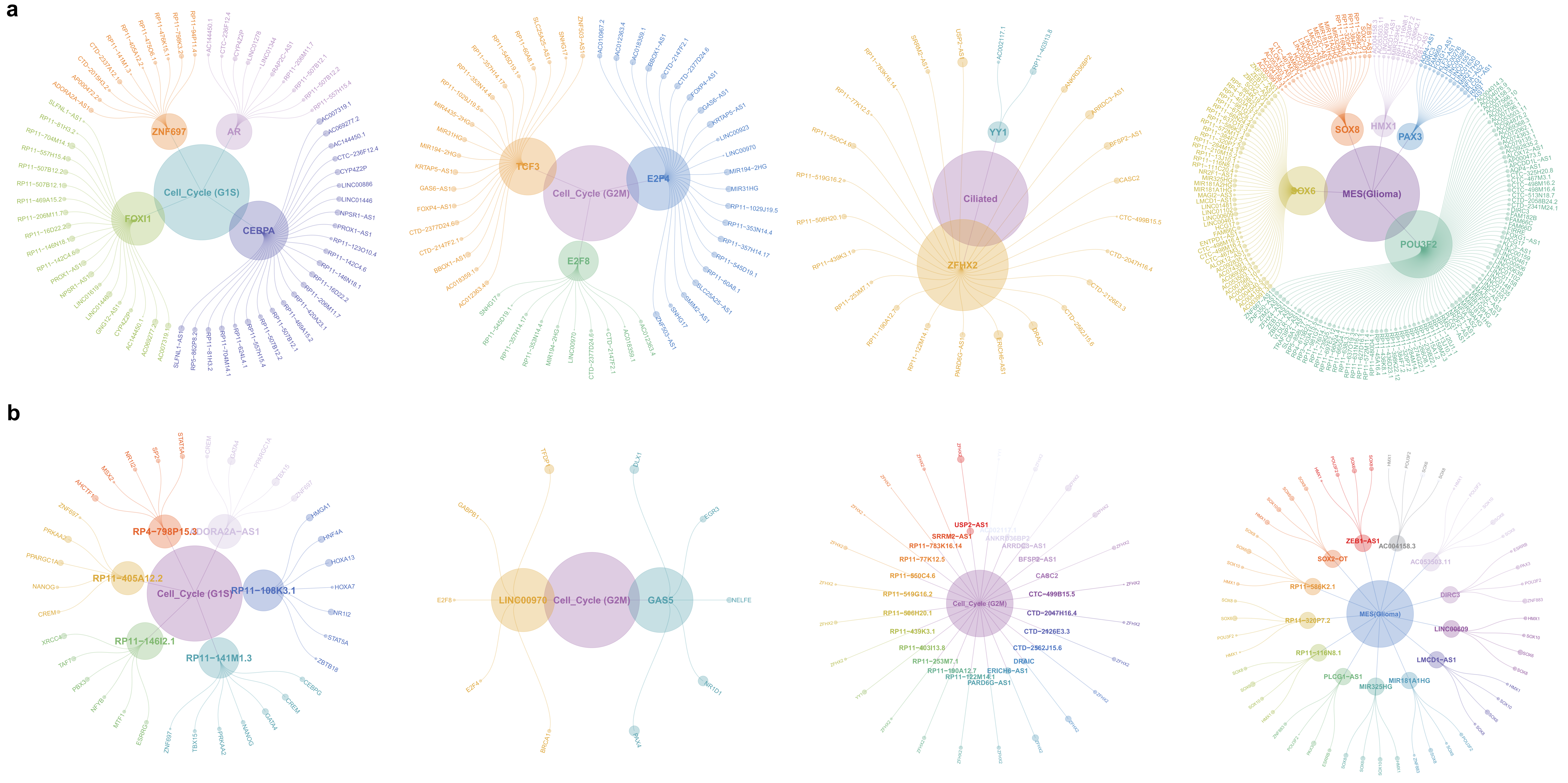
